# A surrogate marker of protection confirms the efficacy of an AddaS03-adjuvanted West Nile virus subunit vaccine

**DOI:** 10.64898/2026.04.20.719748

**Authors:** Atsuko Inoue, Shinji Saito, Kaito Maeda, Yukari Itakura, Shintaro Kobayashi, Michihito Sasaki, Gabriel Gonzalez, William W. Hall, Katsumi Maenaka, Yasuko Orba, Hirofumi Sawa, Koshiro Tabata

## Abstract

West Nile virus (WNV) is the causative agent of fatal West Nile encephalitis. To date, no human vaccine against WNV has been approved. Adjuvants are important for developing effective and affordable vaccines that enhance the immunogenicity and decrease the required antigen doses. In this study, we assessed the efficacy of AddaS03, a synthetic adjuvant analogous to AS03, in a WNV subunit vaccine composed of soluble recombinant envelope protein (sEnv). Using a passive immunization mouse model, we defined the neutralizing antibody titer threshold required for protection against lethal WNV infections and applied this threshold as a surrogate marker to evaluate adjuvant efficacy. AddaS03-adjuvanted formulations elicited markedly higher neutralizing antibody titers compared to Alhydrogel adjuvant 2% (Alhydrogel), even at suboptimal antigen doses, and consistently exceeded the defined protective threshold titer. Moreover, in a sequential challenge mouse model, AddaS03-adjuvanted vaccines completely protected mice from symptomatic WNV infections, whereas Alhydrogel-adjuvanted vaccines failed to confer full protection. Collectively, these findings demonstrate that AddaS03 is a promising adjuvant for WNV subunit vaccine development and highlights the utility of a passive immunization model for defining protective antibody thresholds as a surrogate marker for vaccine evaluation.

## Introduction

West Nile virus (WNV), a member of the *Orthoflavivirus* genus, is a zoonotic virus transmitted from avian hosts to humans, horses and other wildlife species *via* mosquito vectors (1). Approximately 20% of individuals infected with WNV develop clinical symptoms, ranging from a nonspecific febrile illness to severe neuroinvasive diseases such as encephalitis, meningitis, or anterior myelitis that can lead to acute flaccid paralysis (2). Furthermore, approximately 40% of symptomatic individuals are reported to experience long-term sequelae, including fatigue, depression, weakness, and neck or back pain (3). Between 2012 and 2021, there were 3,632 clinical cases in the European Union/European Economic Area countries and 24,029 cases in the United States (4, 5). To date, WNV has been identified in the United States, Europe, West Asia, Africa, and other regions, causing recurrent endemics (6). Hence, WNV infection is expected to pose long-term health and economic burdens worldwide. Thus, the development of an effective human vaccine is urgently required.

Neutralizing antibodies serve as the primary protective agents against orthoflavivirus infections (7). These antibodies recognize viral structural proteins, particularly the envelope protein, which contains the receptor-binding domain and the membrane-fusion domain (8–10). Therefore, previous studies have predominantly focused on vaccine development using soluble recombinant envelope protein (sEnv) to induce neutralizing antibody responses (11, 12). Subunit vaccines incorporating the sEnv have also been evaluated for the prevention of orthoflavivirus infections (13, 14). In a previous study, WNV-derived sEnv formulated with aluminum-based adjuvant (Alum) elicited protective antibody responses in C57BL/6 mice against lethal WNV infection (15). These findings suggest that Alum-adjuvanted sEnv could be a promising vaccine candidate for human use, given that Alum has already been approved for clinical use and is widely used as a vaccine adjuvant with various viral proteins (16). In general, adjuvants are used to enhance vaccine immunogenicity and reduce antigen doses. Although Alum is safe and well-tolerated, relatively high antigen doses are often required to elicit sufficient immune responses (17). Since subunit vaccines are expensive to manufacture, it is critical to minimize the antigen dose to reduce production costs.

To date, various adjuvants have been approved for use in human vaccines such as squalene-based (e.g., AS03 and MF59), nucleic acid-based (e.g., CpG1018), and Lipid-based (e.g., AS01) adjuvants (16). Squalene-based adjuvants AS03 and MF59 exhibit potent immunostimulatory activity by enhancing both humoral and cellular immune responses (18, 19). These adjuvants are already in clinical use in pandemic H5 influenza vaccines (20, 21). Furthermore, AS03 has been reported to enhance the immune response to a lower dose of split influenza vaccine compared to Alum (22). These findings suggest that AS03 is a promising adjuvant for the development of next-generation subunit vaccines against currently untreatable infectious diseases, including WNV infection. However, since adjuvant activity has been reported to vary depending on antigen type (23), it is essential to select the appropriate combination of adjuvant and antigen. To develop a human WNV vaccine, we evaluated the potential of a novel combination of WNV-sEnv and the AS03 adjuvant analogue AddaS03 using a mouse model, based on the intensity of the neutralizing antibody response.

## 2. Materials and methods

### 2.1. Cells

*Aedes albopictus*-derived C6/36 mosquito cells (ATCC, Manassas, VA, USA) were cultured in Eagle’s Minimum Essential Medium (EMEM, Fujifilm Wako Pure Chemical Corporation, Osaka, Japan) supplemented with 10% fetal bovine serum (FBS) and non-essential amino acids (Gibco, Thermo Fisher Scientific, Waltham, MA, USA) at 28°C and 5% CO_2_. Expi293F GnTI(−) cells (Gibco) were cultured in Expi293 expression medium (Gibco) at 37°C and 8% CO□. ExpiSF9 cells (Gibco) were cultured at 27°C in ExpiSf CD medium (Gibco). BTI-TN-5B1-4 (High Five) cells (Gibco) were cultured in Express Five SFM medium (Gibco), supplemented with 2 mM L-glutamine (Fujifilm Wako Pure Chemical Corporation) at 27°C. HEK293T cells (RIKEN BRC, Ibaraki, Japan) were cultured in high-glucose Dulbecco’s Modified Eagle Medium (DMEM, Fujifilm Wako Pure Chemical Corporation) supplemented with 10% FBS at 37°C and 5% CO□. Vero E6 cells (CRL-1586, ATCC, Manassas, VA, USA) were cultured in low-glucose DMEM (Fujifilm Wako Pure Chemical Corporation) supplemented with 10% FBS at 37°C and 5% CO□.

### 2.2. Virus

WNV strain NY-99 6-LP (accession no. AB185914) was inoculated into C6/36 cells. The supernatants were harvested at 5 days post-infection (dpi) and centrifuged at 400 × *g* for 5 minutes to remove cellular debris, and then the virus titer in the supernatants were measured by the TCID_50_ assay as previously described (24). The supernatants were stored at –80°C until use.

### 2.3. Expression and purification of recombinant proteins

For production of rSVP, pCXSN vector encoding precursor membrane, membrane, and envelope proteins (accession no. LC770216) was constructed as previously described (25). Expi293F GnTI(−) cells were transfected with the plasmid using ExpiFectamin 293 Transfection kits (Gibco) following the manufacturer’s protocol. At 5 days post-transfection (dpt), supernatant was harvested and centrifuged at 400 × *g* for 5 minutes to remove cellular debris. The clarified supernatants were filtered with a 0.45 µm polyethersulfone (PES) membranes (Thermo Fisher Scientific). The supernatants were precipitated by ultracentrifugation at ultracentrifugation with a 20% sucrose cushion at 137,788 × *g* for 2 hours using Himac Centrifuge CP100NX (Eppendorf, Hamburg, Germany), and then the pellet was resuspended with DMEM. The rSVP was stored at –80°C until use.

For production of recombinant nonstructural protein 1 (rNS1), the coding sequence for the C-terminus of envelope protein (residues 478 to 501) and NS1 (residues 1 to 352) was optimized for human codon usage (GeneArt, Thermo Fisher Scientific, Regensburg, Germany). The gene was cloned into pCXSN vector with a six-histidine tag at the C-terminus. Expi293F GnTI(−) and cells were transfected with the plasmid following the manufacturer’s protocol. At 7 dpt, supernatants were harvested, centrifuged at 2,000 × *g* for 10 minutes, and purified using a Ni-NTA resin (Cytiva, Marlborough, MA, USA) following the manufacturer’s protocol. Purified rNS1 in elution buffer were concentrated and exchanged into phosphate-buffered saline (PBS) using Amicon Ultra 30 K (Millipore, Burlington, MA, USA). The rNS1 was stored at –80°C until use.

For the production of sEnv, pFastBac vector (Gibco) encoding the ectodomain of the envelope protein (residues 1 to 400) with honeybee melittin signal sequence at the N-terminus and a six-histidine tag at the C-terminus was constructed. MAX Efficiency DH10Bac competent cells (Gibco) were transformed with the pFastBac vector, and then Bacmid DNA encoding baculovirus genome carrying sEnv was extracted. The Bacmid DNA transfected into ExpiSf9 cells at 2.5×10^6^ cells/mL using ExpiFectamine Sf Transfection Reagent (Gibco) following the manufacturer’s protocol. Supernatants containing recombinant baculoviruses were harvested after 7 dpi and centrifuged at 2,000 × *g* for 10 minutes to remove cellular debris. The supernatant was inoculated into High Five cells at 1.0×10^6^ cells/mL and then the cells were maintained for 5 days. Following the incubation, the supernatants were processed in the same manner as for rNS1 production. Further purification was carried out by size exclusion chromatography using a Superdex 200 Increase 10/300 GL column (Cytiva) on an ÄKTA go system (Cytiva). The sEnv was stored at –80°C until use.

### 2.4. Ethics Statement and Biosafety

All animal experiments were conducted in accordance with the regulations on animal experiments at the National University Corporation, Hokkaido University Regulations on Animal Experimentation. The experimental design was reviewed and approved by the Institutional Animal Care and Use Committee of Hokkaido University (approval no. 24-0034). All experiments with live virus were conducted in a biosafety level 3 (BSL3) laboratory.

### 2.5. WNV infection in mice

Female BALB/c mice (10-week-old, n=6 or 12) were intraperitoneally inoculated with 10^2^, 10^3^, 10^4^, or 10^5^ TCID_50_ /100 µL of WNV. Mice were monitored daily for body weights and survival. Based on humane endpoints, mice showing body weight loss of > 20% were euthanized with an overdose of isoflurane. At 13 dpi, the surviving mice were euthanized with an overdose of isoflurane, and the blood was collected to obtain convalescent serum. Serum samples were heat-inactivated at 56°C for 30 minutes and stored at -80°C until use.

### 2.6. Evaluation of antibody responses

Antibody responses were evaluated using an indirect Enzyme-linked immunosorbent assay (ELISA). rNS1 was coated overnight at 4°C on 96-well half-area plates (Corning, Tewksbury, MA, USA), and SVPs were coated overnight at 4°C on 384-well plates (Corning). Nonspecific binding was blocked by incubating the plates with 5% FBS in PBS for 30 minutes at room temperature. Then, diluted mouse serum samples and pooled convalescent serum (used for calibration curve) were added to the plates and incubated at 37°C for 2 hours. After washing six times with PBS containing 0.01% Tween 20 (PBST), the samples were incubated with horseradish peroxidase (HRP)-conjugated anti-mouse IgG (Sigma-Aldrich) at 37°C for 1 hour. Following another wash, 1-Step Ultra TMB-ELISA Substrate Solution (Thermo Fisher Scientific) was added. The reaction was then stopped by adding 2M sulfuric acid. The absorbances at 450 nm and 690 nm were measured using an Infinite 200 PRO (Tecan, Männedorf, Switzerland), and the optical density at 450 nm (OD_450nm_) value was calculated as the difference between the absorbances at 690 nm and 450 nm. Antibody titers of an individual serum sample were expressed as ELISA units, determined by a calibration curve generated from serial dilutions of pooled convalescent serum. The ELISA unit of the pooled convalescent serum was defined as the reciprocal of the highest dilution factor exceeding the cutoff value (mean plus three times standard deviations (SD) of the naive serum) (26).

### 2.7. Neutralization assays

Microneutralization assays using WNV-derived single-round infectious particles (WNV-SRIPs) were conducted with slight modifications of the previously reported procedures (27). Briefly, WNV-SRIPs were generated by co-transfection into HEK293T cells with two plasmids: pC-CprME, encoding the WNV-derived capsid, precursor membrane, membrane, and envelope proteins, and pCMV-WNVrep-EGFP, encoding a replicon containing the full-length WNV genome with structural genes replaced by enhanced green fluorescent protein (EGFP). The culture media were replaced with fresh medium at 12 hours post-transfection, and the cells were subsequently incubated at 28°C. Supernatants were collected at 4 to 6 dpt and centrifuged at 400 × *g* for 5 minutes to remove cellular debris. For titration of the WNV-SRIPs, Vero E6 cells at 1.5×10^5^ cells/mL were seeded in 96-well plates (Corning). The supernatants were initially diluted 1:50, followed by 2-fold serial dilutions and inoculated into each well. Three days after incubation at 37°C, snapshots of each well were obtained using an IN Cell Analyzer 2500HS (GE Healthca0re, Chicago, IL, USA), and EGFP-positive cell numbers were counted using Fiji/ImageJ (28). The titers were expressed as infectious units (IU) per mL.

For neutralization assays, Vero E6 cells at 1.5×10^5^ cells/mL were seeded in 96-well plates. Two-fold serial diluted sera were mixed with 300 IU of the WNV-SRIPs at a 1:1 ratio. After incubation for 2 hours at 37°C, the cells were inoculated with the serum-SRIPs complexes. Three days post-incubation, EGFP-positive cell were analyzed as described above. Neutralization (NT) titers were defined as the reciprocal of the highest serum dilution that resulted in a 50% reduction in EGFP-positive cell counts compared to WNV-SRIPs control.

### 2.8. WNV challenge in passively immunized mice

A serum sample with a 128,000 NT titer /mL was prepared by pooling portions of the convalescent serum listed in Supplementary Table 1 (#16, #24, #32, and #37). This pooled serum was serially diluted from 1:1 to 1:1,000. Female BALB/c mice (5-week-old, n=4) were intravenously injected with 50 µL of each diluted serum sample or PBS used as a negative control, then the mice were challenged with 100 TCID_50_ /100 µL of WNV *via* intraperitoneal inoculation at 24 hours after injection of serum or PBS. Their body weights and survival were monitored daily for the following 14 days. Mice were euthanized with an overdose of isoflurane when their body weights reached humane endpoints as described above. At 14 dpi, surviving mice were euthanized with an overdose of isoflurane, and blood was collected and separated into serum by centrifugation. Serum samples were heat-inactivated at 56°C for 30 minutes and stored at –80°C until use.

### 2.9. Assessment of serum dilution rate *in vivo*

The pre-transfer serum samples were prepared by pooling the convalescent serum listed in Supplementary Table 1 (#27, #29, and #40). The pre-transfer serum samples were divided into four serum samples. Each serum sample was diluted 1:10 (n=2) or 1:100 (n=2) and passively transferred intravenously into five-week-old mice. At 24 hours post-transfer, mice were euthanized with an overdose of isoflurane, and blood was collected and processed to obtain heat-inactivated serum (post-transfer serum). Pre- and post-transfer serum samples were stored at –80°C until use. Neutralization assay and ELISA were conducted using each serum samples to assess the dilution rate in the mice body after passive transfer of serum.

### 2.10. SDS-PAGE and immunoblotting (IB)

Purified sEnv was separated by sodium dodecyl sulfate-polyacrylamide gel electrophoresis (SDS-PAGE). One sample (2 µg /lane) was stained with Coomassie brilliant blue (CBB). Another (0.5 µg /lane) was transferred to polyvinylidene difluoride (PVDF) membranes (Millipore). The membranes were blocked with 5% skimmed milk in PBST and incubated with pan-orthoflavivirus anti-fusion loop monoclonal antibody 4G2 (supernatant from D1-4G2-4-15 cells, ATCC) at 4°C overnight. After washing with PBST three times, HRP-conjugated anti-mouse IgG antibody (Sigma-Aldrich) was used as the secondary antibody. Signals were visualized using Immobilon Western Chemiluminescent HRP substrate (Millipore) and LuminoGraph II EM (ATTO, Tokyo, Japan).

### 2.11. Vaccination of mice

Female BALB/c mice (5-week-old, n=6) were subcutaneously injected with sEnv (0, 0.1, 0.5, 1, or 5 µg/body), either alone or in combination with 50% v/v AddaS03 (InvivoGen, San Diego, CA, USA) or 50% v/v Alhydrogel (Alhydrogel adjuvant 2%; InvivoGen). At 13 days after the primary vaccination, blood samples were collected from the orbital sinus of mice anesthetized with isoflurane. At 2 weeks after the priming vaccination, the mice were boosted with the same vaccine formulations. At 3 weeks after the boost vaccination, mice were euthanized using an overdose of isoflurane, and blood samples were collected and serum collected. Serum samples were heat-inactivated at 56°C for 30 minutes and stored at –80°C until use.

### 2.12. Sequential WNV challenge in vaccinated mice

Female BALB/c mice (5-week-old, n=4) were subcutaneously vaccinated with sEnv (1 or 5 µg/body) in combination with 50% v/v AddaS03 or Alhydrogel. Vaccination schedules described in Section 2.11. At 3 weeks after boost vaccination, the mice were challenged with 3,000 TCID_50_/100 µL of WNV *via* the intraperitoneal route, and their body weights were monitored daily for the following 14 days. Mice were euthanized with an overdose of isoflurane upon reaching the humane endpoint. At 14 dpi, surviving mice were euthanized with an overdose of isoflurane, and blood was collected and processed to obtain heat-inactivated serum. Serum samples were stored at –80°C until use.

### 2.13. Statistical analyses

All statistical analyses were performed using GraphPad Prism software version 8.0 (Dotmatics, Boston, MA, USA). Detailed analysis is described in each figure legend.

## 3. Results

### 3.1. A passive immunization model reveals the minimum neutralization titer required for protection against WNV

Previous studies have demonstrated that passive immunization, achieved by transferring sera from vaccinated mice into naïve 5-week-old recipients, can be used to evaluate neutralizing antibody responses that confer protection against lethal WNV infection (29). The passive immunization model was employed to define the serum NT titer required for protection against lethal WNV infection. In this model, 5-week-old mice were intravenously administered serum samples with pre-determined NT titers and subsequently challenged with WNV at 24 hours after transfer. (**Fig. 1A**). To prepare serum for use in this model, NT titer was first evaluated in pooled convalescent sera collected from mice that had survived WNV infection (**Fig. 1B**). The pooled serum, which had 128,000 NT titer per mL, was serially diluted 10-fold, and 50 µL of each diluted serum was passively transferred into naïve 5-week-old mice. After infection with 100 TCID_50_ of WNV, body weight change and survival were monitored daily. At 6 to 7 dpi, vehicle-treated mice exhibited rapid body weight loss, and three of four mice succumbed to the infection (**Figs. 1C, 1D, and S1A**). Notably, passive transfer of undiluted or 10-fold diluted serum completely protected naïve mice from symptomatic or lethal infection. In contrast, transfer of 100-fold- or 1,000-fold-diluted serum conferred only partial or no protection, respectively (**Figs. 1C, 1D, and S1A**). To further characterize the protective effect, antibody titers against the nonstructural protein 1 (NS1) were measured by ELISA using convalescent sera collected from the passively transferred mice at 14 dpi. The antibodies to NS1 are induced by orthoflavivirus infections due to their presence in both the intracellular and extracellular compartments (30, 31). The ELISA showed that anti-NS1 antibody titers in mice receiving 100- or 1,000-fold diluted sera were comparable to those in vehicle-treated mice (**Fig. S1B**). In contrast, mice passively immunized with undiluted or 10-fold diluted sera exhibited a 21.9-fold or 4-fold reduction in anti-NS1 antibody titers, respectively, compared to the control group. These NS1-specific antibody profiles are consistent with the protective effects observed in mice treated with undiluted or 10-fold diluted sera against WNV infection (**Figs. 1C and 1D**).

**Figure 1.**
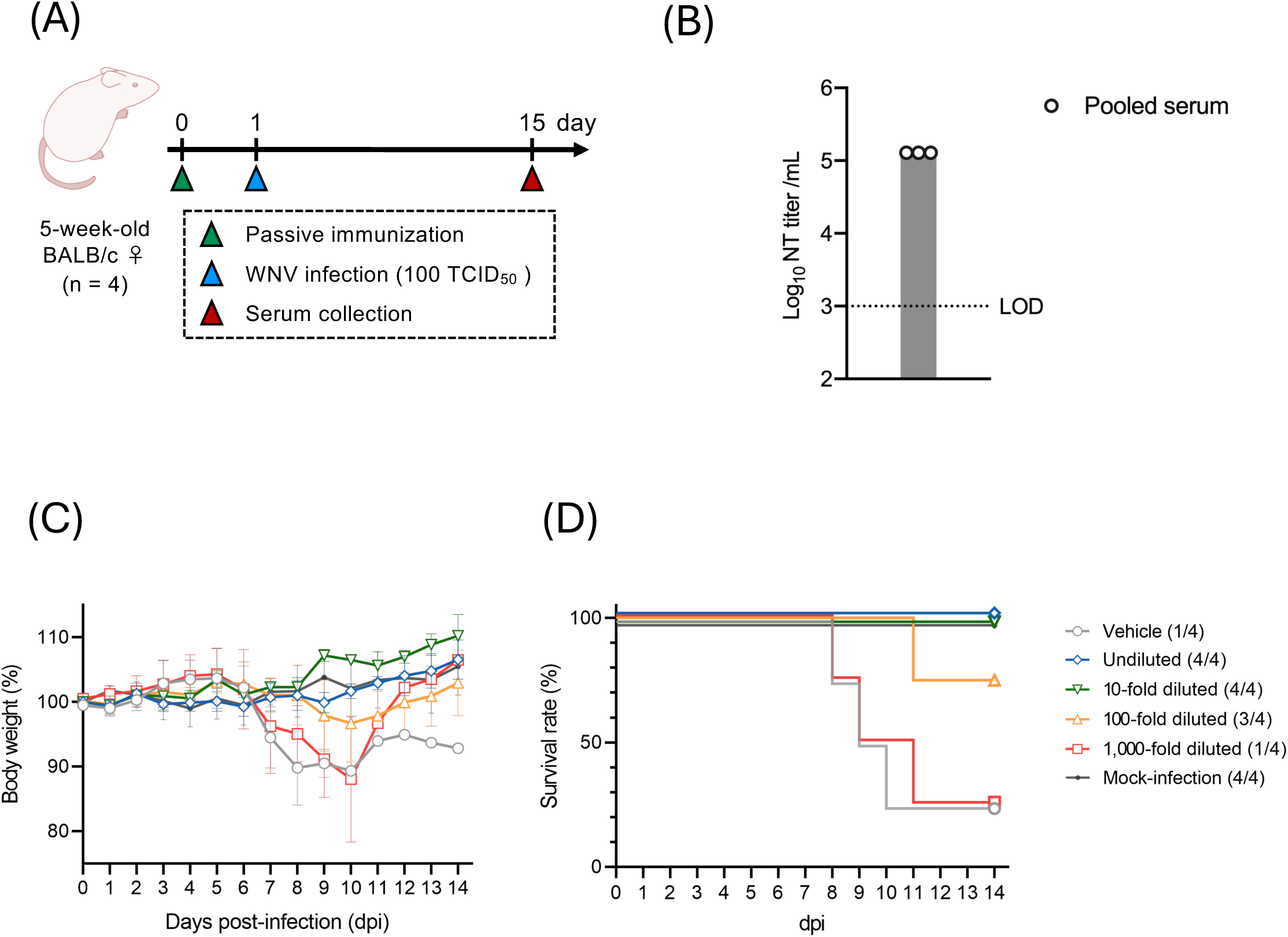
Passive transferred serum protected lethal WNV infection in mice(5-week-old). (**A**) Schematic illustration of passive transfer model of serum to define the protective neutralizing antibody titer against lethal WNV infection. Pooled serum was passively transferred to naïve five-week-old mice at 24 hours before challenge with WNV. (**B**) Neutralization titer of pooled convalescent serum from WNV infection was determined. The dotted line indicates the LOD. (**C and D**) BALB/c mice (5-week-old, n=4) were infected with 100 TCID_50_ of WNV and monitored daily for body weights (**C**) and survival rate (**D**) for 14 days. The numbers shown in parentheses in the legend represent the number of surviving mice out of the total. Data is represented as the mean±SD of technical replicates (n=3) (**B**) and biological replicates (n=4) (**C**).

To estimate the NT titer required for protection against lethal WNV infection, the *in vivo* dilution rate of passively transferred serum was calculated. A neutralization assay was conducted using the serum sample prior to transfer compared as sera collected from recipient mice 24 hours after passive transfer. The NT titer in the post-transfer serum was below 1,000 NT titer/mL, the limit of detection (LOD), whereas binding antibody titers were detected in both serum samples (**Figs. S2A and S2B**). Given the strong correlation between binding and neutralizing activities in the pre-transferred serum (r=0.9014, R^2^=0.8125), the dilution factor was estimated based on changes in the binding antibody titers of pre- and post-transfer sera (**Figs. S2C, S2D, and S2E**). Based on the binding antibody titers, the *in vivo* dilution rate was estimated to be approximately 41.15-fold (**Fig. S2F**). The passive transfer of 10-fold diluted serum (12,800 NT titer/mL) conferred complete protection against lethal WNV infection (**Figs. 1C and 1D**). Considering the estimated 41.15-fold *in vivo* dilution, the NT titer of the 10-fold diluted serum in recipient mice was approximately 311 NT titer/mL. Thus, the NT titer required in serum to confer protection against the WNV infection was defined as 311 NT titer/mL. In subsequent experiments, threshold NT titer (≧311 NT units/mL) was used as a surrogate marker to evaluate the efficacy of adjuvants in vaccine formulations.

### 3.2. The AddaS03-adjuvanted subunit vaccine induced neutralizing antibody responses that exceeded the threshold required for protection

In this study, AddaS03, a formulation similar to the approved adjuvant AS03, was assessed for its adjuvanticity in combination with a subunit vaccine, based on the defined threshold (≧311 NT units/mL). WNV-derived sEnv, produced using the baculovirus-insect cell expression system, was used as a vaccine antigen (**Fig. 2A**). BALB/c mice (5-week-old) were immunized twice at two-week intervals using a vaccine formulation containing varying doses of sEnv antigen and either Alhydrogel adjuvant 2% (Alhydrogel) or AddaS03. Blood samples were collected from each mouse at 13 days after the priming vaccination and 3 weeks (at 35 days after the primary vaccination) after the boosting vaccination (**Fig. 2B**).

**Figure 2.**
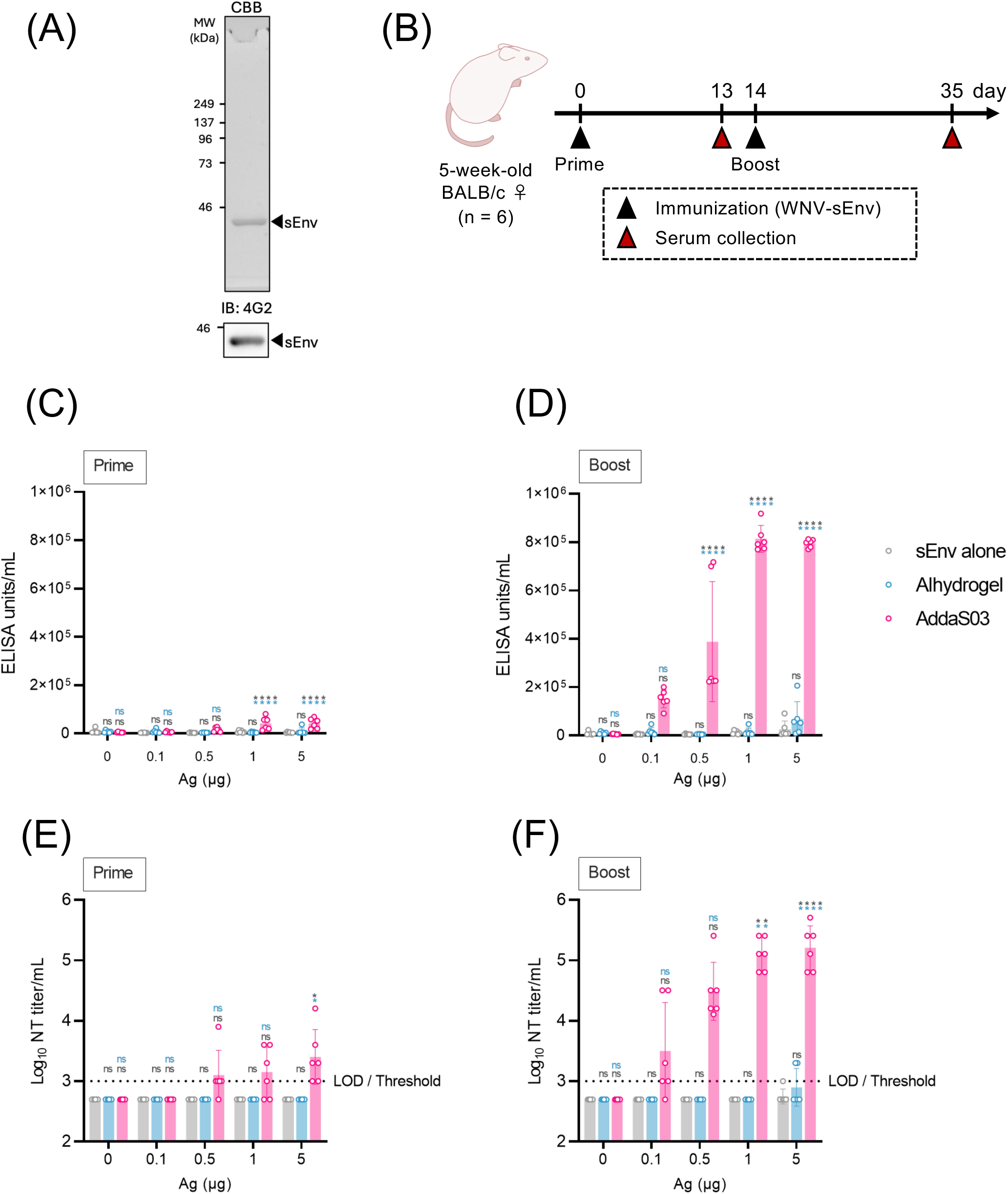
Neutralizing antibody titers in mice immunized with AddaS03□adjuvanted sEnv exceeded the threshold titers and were higher than those observed in the other groups. (**A**) Identification of purified sEnv by CBB staining and immunoblotting using a pan-orthoflavivirus anti-fusion loop monoclonal antibody (clone 4G2). (**B**) Five-week-old BALB/c mice (5-week-old, n=6) were immunized subcutaneously with a vaccine formulation containing various antigen doses (0, 0.1, 0.5, 1, or 5 μg) with or without adjuvant (Alhydrogel or AddaS03) twice at 2 weeks interval. Serum samples were collected at 13 and 35 days after the first immunization. (**C to F**) Binding and neutralizing antibody titers in murine sera after priming (**C and E**) and boosting immunizations (**D and F**) were determined by indirect ELISA using WNV SVP and by a neutralization assay, respectively. Dotted line indicates the LOD. The threshold titer estimated as sufficient to protect against lethal WNV infection in Figure 1 is also indicated as the LOD, since the threshold value was below the LOD. Data are represented as the mean±SD of biological replicates (n=6). Statistical analysis was performed using two-way ANOVA followed by Dunnett’s multiple comparison test. All statistical differences are determined in comparison with the sEnv alone (gray) or Alhydrogel (blue) group within the same antigen dose (\**p*<0.01, \*\**p*<0.005, \*\*\*\**p*<0.0005). ns, not significant.

To evaluate antibody responses to sEnv, an indirect ELISA using WNV subviral particles (SVPs), composed of the precursor membrane and envelope protein, was performed on serum samples collected after priming and boosting. The ELISA showed no significant difference in antibody titers across different antigen doses between sEnv alone and sEnv with Alhydrogel. In contrast, priming with the AddaS03-adjuvanted vaccine induced significantly higher antibody titers than sEnv at both 1□µg and 5□µg doses with or without Alhydrogel (**Fig. 2C**). Notably, sera collected after boosting with the AddaS03-adjuvanted vaccine showed a marked increase in the binding antibody titers, especially in groups that received an antigen dose of 0.5□μg or higher, which exhibited significantly higher titers compared to the sEnv alone and Alhydrogel-adjuvanted groups (**Fig. 2D**).

Neutralizing activity against WNV was further assessed using a neutralization assay with WNV-derived single-round infectious particles (WNV-SRIPs). As already described, the NT titer required for protection was estimated to be 311 NT titer/mL. However, since this value was below the LOD (1,000 NT titer/mL), the LOD was adopted as the protective threshold in this experiment. Surprisingly, in the AddaS03-adjuvanted vaccine groups, NT titers exceeding the threshold were already induced after priming with the highest antigen dose (**Fig. 2E**). Furthermore, following boosting, even the group receiving the lowest antigen dose produced neutralizing antibody with NT titers exceeding the threshold, and significantly higher NT titers were observed in the 1□µg and 5□µg antigen dose groups compared to the other groups (**Fig. 2F**). Sera from some mice immunized with the highest antigen dose, either with sEnv alone or sEnv with Alhydrogel, exceeded the threshold NT titer following boosting (**Fig. 2F**).

### 3.3. Body weight change serves as a surrogate marker for the establishment of WNV infection

To evaluate vaccine efficacy against WNV infection using a sequential challenge mouse model, we assessed the susceptibility of BALB/c mice (10-week-old) to WNV infection. The mice were infected intraperitoneally route with 10^2^ to 10^5^ TCID_50_ of WNV, and survival and body weight changes were monitored daily. In the groups infected with 10^2^, 10^4^, and 10^5^ TCID_50_ of WNV, a subset of mice exhibited body weight loss, whereas the remaining mice showed no apparent changes (maximum weight loss >10%) were observed in the other mice. (**Fig. S3A**). All mice infected with 10^3^ TCID_50_ of WNV exhibited body weight loss. However, no differences in mortality rates were observed among the groups (**Fig. S3B**). To identify differences in mice surviving WNV infection, the symptomatic outcomes, including maximum body weight loss rate and mortality, were summarized (**Supplementary Table1**). It was observed the WNV infection at a 10^3^ TCID_50_ dose induced notable symptoms, including mortality or >10% body weight loss, in all 10-week-old mice. Conversely, some surviving mice infected at the other doses (10^2^, 10^4^, or 10^5^ TCID_50_) exhibited maximum body weight loss below 10%. These findings suggest that 10-week-old mice are relatively resistant to lethal WNV infection. Nevertheless, body weight loss in mice infected with 10^3^ TCID_50_ served as a consistent surrogate marker of successful infection.

### 3.4. The AddaS03-adjuvanted subunit vaccine completely protected against symptomatic WNV infection

To validate the protective efficacy of AddaS03 against symptomatic WNV infection, a sequential infection model utilizing 3,000 TCID_50_ of WNV was employed, as 10^3^ TCID_50_ had previously been shown to induce >10% body weight loss in all mice (Section 3.3). The antigen dose used for vaccination was determined according to the significant differences observed in neutralizing antibody responses between sEnv antigens formulated with either Alhydrogel or AddaS03 (**Fig. 2F**). BALB/c mice (10-week-old) immunized twice with either 1 or 5 µg of sEnv antigen in combination with Alhydrogel or AddaS03 were subsequently challenged with WNV (**Fig. 3A**). Following WNV infection, consistent body weight loss was observed in all vehicle-immunized mice (**Fig. S4**). Although mortality occurred in both Alhydrogel-adjuvanted groups, some mice showed no obvious signs of weight loss (**Figs. 3B, 3C, and S4**). Moreover, in the group received the 1□µg antigen dose, despite NT titers remained below the threshold NT titer (**Fig. 2F**), body weight loss was not observed in one of four mice (**Figs. 3B and S4**). These findings suggest that the minimum protective NT titer may fall below the LOD. Importantly, mice immunized with the AddaS03-adjuvanted vaccine demonstrated complete protection against symptomatic WNV infection at both antigen doses, without exhibiting body weight loss (**Figs. 3B, 3C and S4**).

**Figure 3.**
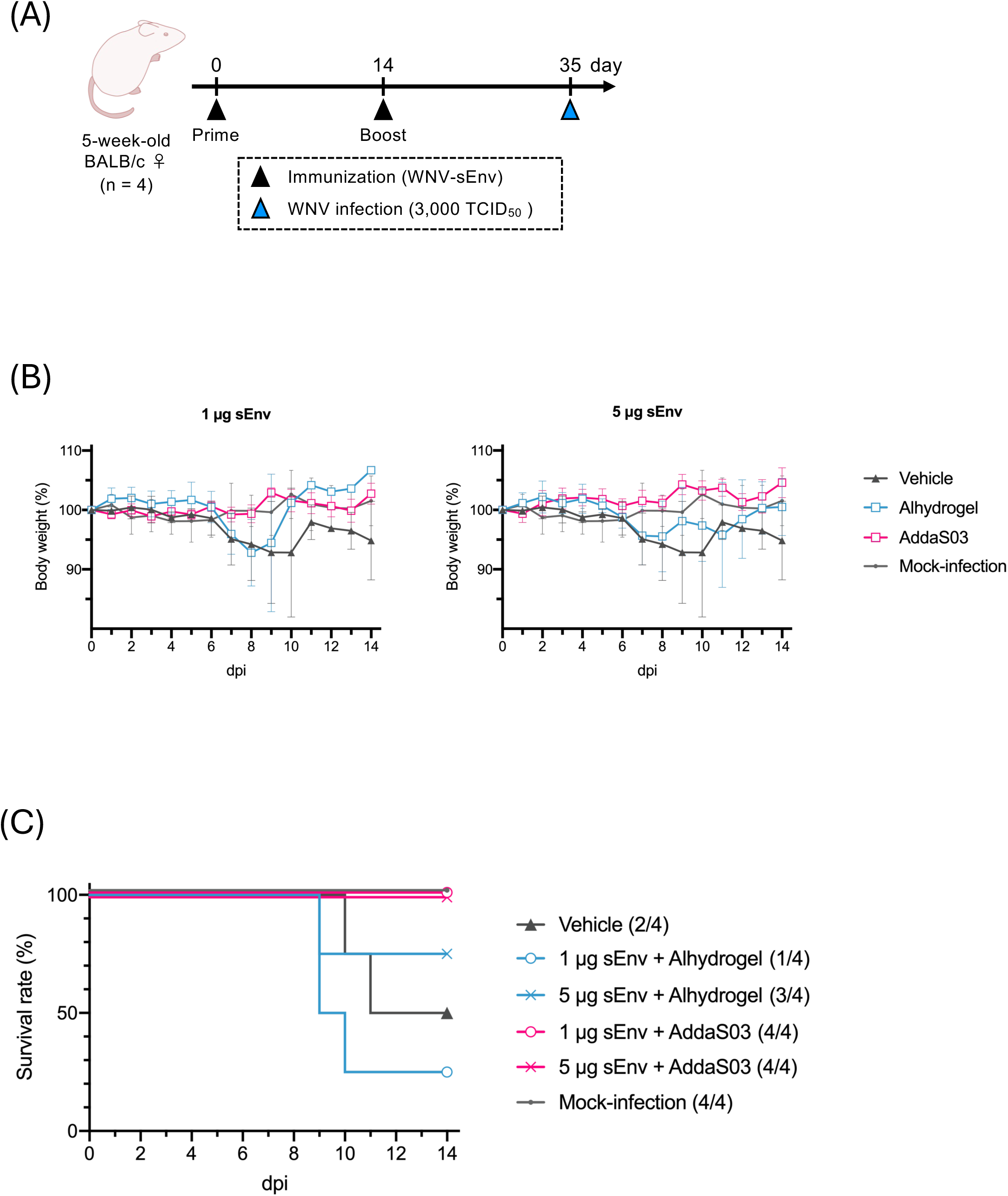
Vaccination with AddaS03-adjuvanted sEnv provided the mice with protective immunity against symptomatic WNV infection. (**A**) BALB/c mice (5-week-old, n=4) were immunized subcutaneously with a vaccine formulation containing 1 or 5 µg of WNV-sEnv with adjuvant (Alhydrogel or AddaS03) or PBS (Vehicle) twice at 2 weeks interval. On 35 days after the first immunization, the mice were challenged with 3,000 TCID_50_ of WNV. (**B and C**) The infected mice were monitored daily for body weights (**B**) and survival (**C**) for 14 days after the challenge. Data in the graph are expressed as the mean±SD of body weight changes.

## 4. Discussion

In this study, we compared the adjuvant effects of the well-validated Alhydrogel or AddaS03 under low-antigen-dose conditions on a subunit vaccine based on WNV-sEnv to develop a novel subunit vaccine formulation against WNV infection. Our results demonstrated that the subunit vaccine combined with AddaS03 induced NT titers markedly exceeding the threshold required for protection, even at a sEnv dose of 0.5 µg (**Fig. 2F**). Furthermore, the AddaS03-adjuvanted vaccine protected mice from WNV infection, which was accompanied by symptoms, including mortality and body weight loss, whereas the Alhydrogel-adjuvanted vaccine failed to provide full protection for all mice in the group (**Figs. 3B and 3C**). Consistent with our findings, previous studies using different recombinant hemagglutinin and neuraminidase proteins derived from influenza A viruses have also demonstrated that AddaS03 induces more robust and functional immune responses than Alhydrogel adjuvant (32). Taken together, these results suggest that AddaS03 exhibits superior adjuvant activity compared with Alhydrogel in some subunit vaccines, and that this effect may not be dependent on the antigen source. Moreover, AddaS03 is a synthetic formulation based on AS03, an adjuvant approved for clinical use in humans as part of the pandemic H5 influenza vaccine (20). Given this background, AddaS03 is considered safe and its successful application to the development WNV subunit vaccine is anticipated. In addition, WNV subunit vaccines formulated with the clinically approved AS03 adjuvant could be promising candidates for protection against WNV infection.

Significant differences in the adjuvant effects on neutralizing antibody induction were observed between AddaS03 and Alhydrogel. The Alum (the same formulation as Alhydrogel) is generally considered to activate immune responses through relatively limited mechanisms, including cell death at the injection site and the release of host-derived DNA, which leads to a predominant Th2-type response (33). In contrast, AddaS03 (AS03-based) engages multiple pathways, with squalene contributing to innate immune activation and α-tocopherol enhancing cytokine/chemokine production and antigen uptake by antigen-presenting cells (34, 35). Consequently, AddaS03 can promote both Th2-type antibody responses and Th1-type cellular responses (36). While these mechanisms may explain the superior adjuvant activity of AddaS03 in this study, it should be noted that the present study did not directly investigate the underlying immunological pathways. Further studies will therefore be required to clarify the precise mechanisms involved.

Here, a passive immunization mouse model was used to evaluate protective antibody responses against WNV infection, and the threshold for protective antibody titers was determined. This threshold clearly reflected the impact of AddaS03-mediated immune activation on protection against WNV infection (**Figs. 1 and 2**). In the conventional sequential challenge model using the same vaccinated mice, the AddaS03-formulated vaccine completely protected mice from symptomatic WNV infections, whereas the Alum-adjuvanted vaccine showed partial protective effects (**Fig. 3B and 3C**). Collectively, these findings suggest that the passive immunization model employed is useful for evaluating the neutralizing antibody titers required for protection in vaccine development. This approach is particularly valuable for studying vaccine efficacy against other infections such as Orthoflaviviruses, in which susceptibility to infection decreases with increasing age in mice (37). However, this model is primarily suitable for evaluating antibody responses that contribute to protection against infection, and it does not allow assessment of cellular immune responses. This limitation is especially relevant for adjuvants such as AddaS03, which are capable of inducing cellular as well as humoral immunity. Therefore, complementary approaches are required to evaluate cellular responses.

Together, these findings highlight the potential of the AddaS03 as a promising adjuvant for WNV subunit vaccines, offering superior antibody responses and complete protection compared with Alhydrogel. These results may inform the rational design and validation of future vaccines against WNV and related Orthoflavivirus infections.

## Acknowledgements

This work was supported by the Japan Agency for Medical Research and Development (AMED) under grant number JP25wm0325073 (Y.O.), JP24wm0225044 (Y.I.), JP223fa627005 (G.G., Y.I., S.S., H.S., and K.T.), JP21wm0125008 (H.S.), and JP253fa827031 (K.T.), by the Ministry of Education, Culture, Sports, Science and Technology, Japan (MEXT) /Japan Society for the Promotion of Science (JSPS) KAKENHI under the grant number JP23K27831 (Y.O.) and JP25K20567 (K.T.), by the Japan Science and Technology Agency (JST) Moonshot R&D under the grant number JPMJMS2025 (Y.O.), by The Uehara Memorial Foundation (K.T.), and by the Promotion Project for Young Investigators in Hokkaido University (H.S.).

## Author contributions

K.T. conceived and designed the research. A.I., S.S., K.M., and K.T. conducted experiments. A.I. and K.T. analyzed data. A.I. and K.T. wrote the draft. All authors reviewed and contributed to the preparation of the manuscript.

## Competing interests

The authors declare no competing interests.

## Supplementary information

**Supplementary Table 1.** Summary of body weight loss after WNV infection in supplementary Figure 3.

| ID | Infectious dose | Outcome<br>(survived/died) | Maximum weight loss (%)<br>(○: >10%) |  |
| --- | --- | --- | --- | --- |
| 1 | 10<sup>2</sup> TCID<sub>50</sub> | died | 20.0 | ○ |
| 2 |  | died | 20.0 | ○ |
| 3 |  | survived | 9.5 | - |
| 4 |  | died | 20.0 | ○ |
| 5 |  | survived | 7.1 | - |
| 6 |  | died | 18.6 | ○ |
| 7 |  | died | 13.4 | ○ |
| 8 |  | died | 20.0 | ○ |
| 9 |  | died | 20.0 | ○ |
| 10 |  | survived | 12.1 | ○ |
| 11 |  | died | 12.0 | ○ |
| 12 |  | died | 20.0 | ○ |
| 13 | 10<sup>3</sup> TCID<sub>50</sub> | died | 4.4 | - |
| 14 |  | died | 20.0 | ○ |
| 15 |  | died | 14.2 | ○ |
| 16 |  | survived | 14.2 | ○ |
| 17 |  | died | 18.4 | ○ |
| 18 |  | died | 16.4 | ○ |
| 19 |  | died | 20.0 | ○ |
| 20 |  | died | 20.0 | ○ |
| 21 |  | died | 8.8 | - |
| 22 |  | died | 17.8 | ○ |
| 23 |  | died | 13.7 | ○ |
| 24 |  | survived | 15.1 | ○ |
| 25 | 10<sup>4</sup> TCID<sub>50</sub> | died | 19.0 | ○ |
| 26 |  | died | 19.6 | ○ |
| 27 |  | survived | 6.4 | - |
| 28 |  | died | 20.0 | ○ |
| 29 |  | survived | 4.6 | - |
| 30 |  | died | 14.2 | ○ |
| 31 |  | died | 9.4 | - |
| 32 |  | survived | 5.4 | - |
| 33 |  | survived | 4.4 | - |
| 34 |  | died | 20.0 | ○ |
| 35 |  | died | 11.8 | ○ |
| 36 |  | died | 3.2 | - |
| 37 | 10<sup>5</sup> TCID<sub>50</sub> | survived | 5.2 | - |
| 38 |  | died | 5.3 | - |
| 39 |  | died | 19.7 | ○ |
| 40 |  | survived | 2.8 | - |
| 41 |  | died | 10.0 | - |
| 42 |  | died | 13.3 | ○ |

**Figure S1.** (**A**) Individual and mean body weight changes corresponding to those in Figure 1C are shown. The numbers shown in parentheses next to each graph title represent the number of surviving mice out of the total. The thick line represents the mean body weight, and the thin lines represent individual mice. Mean data in the graph are expressed as the mean±SD of surviving mice (n=1–4 / group). (**B**) Binding antibody titer to recombinant NS1 was measured from convalescent serum.

**Figure S2.** (**A and B**) NT titers against WNV-SRIPs (**A**) and binding antibody titers to SVP (**B**) were determined by neutralization tests and indirect ELISA, respectively, using 10-fold-and 100-fold-diluted convalescent sera. The dotted line indicates the LOD. (**C and D**) Binding activity to WNV-SVP (**A**) and neutralizing activity against WNV-derived SRIPs (**B**) were determined using pooled convalescent serum from WNV infected mice. (**E**) Correlation between binding and neutralizing activities was analyzed. The Pearson correlation coefficient (r) is shown in the Figure. (**F**) Serum dilution rate in the mouse body was estimated from the binding antibody titer of pre- and post-transfer sera.

**Figure S3.** BALB/c mice (10-week-old, n=12, 12, 12, or 6) were intraperitoneally inoculated with 10^2^, 10^3^, 10^4^, or 10^5^ TCID_50_ of WNV and monitored daily for body weights (**A**) and survival (**B**) for 13 days. (**A**) The thick line represents the mean body weight, and the thin lines represent individual mice. Mean data are represented as the mean±SD of body weight changes.

**Figure S4.** Individual and mean body weight changes corresponding to those in Figure 3B are shown. The numbers shown in parentheses next to each graph title represent the number of surviving mice from the total. The thick line represents the mean body weight, and the thin lines represent individual mice. Data in the graph are expressed as the mean±SD of surviving mice (n=4 / group).

## Notes

### Competing Interest Statement

The authors have declared no competing interest.

