## Supplementary figures and images for "A surrogate marker of protection confirms the efficacy of an AddaS03-adjuvanted West Nile virus subunit vaccine"

### Supplementary Figure

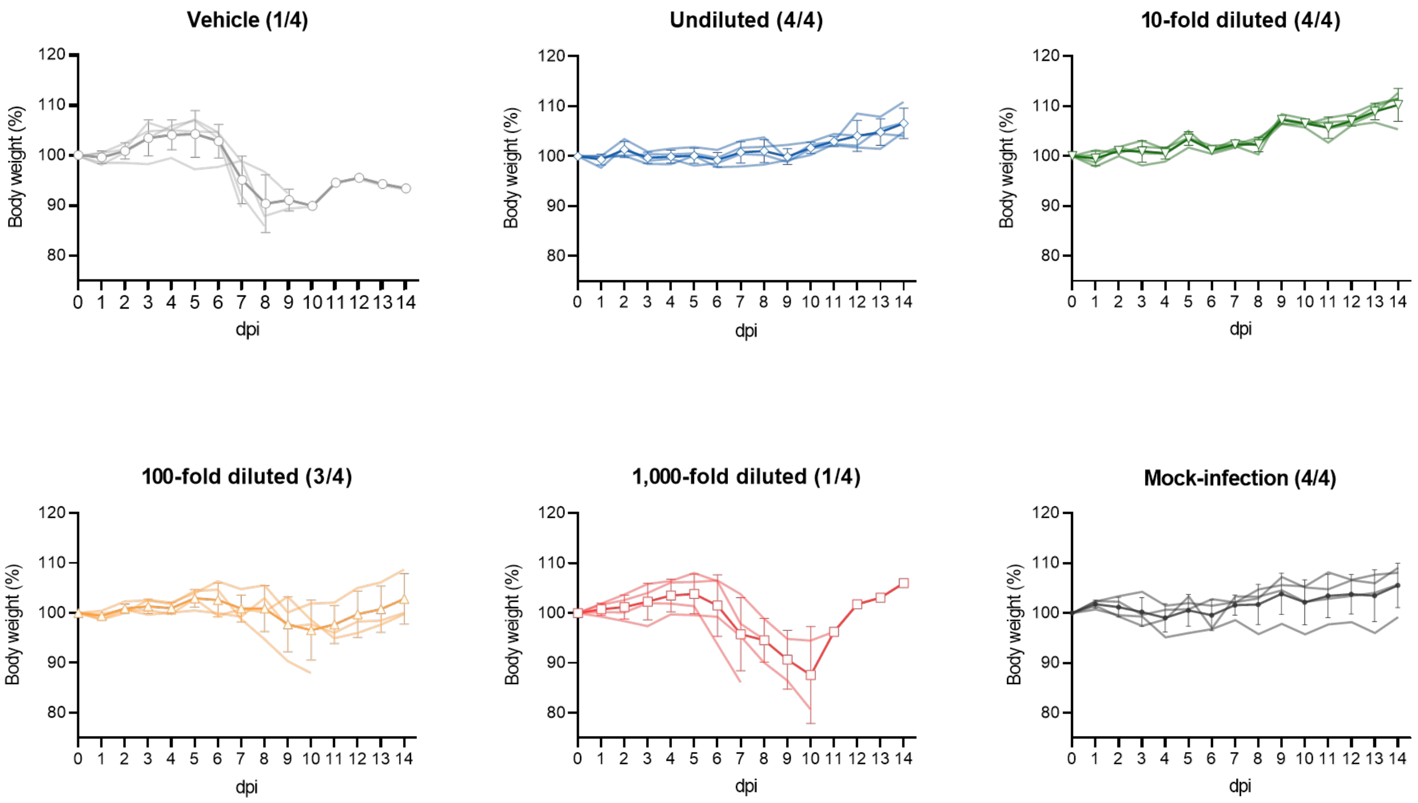

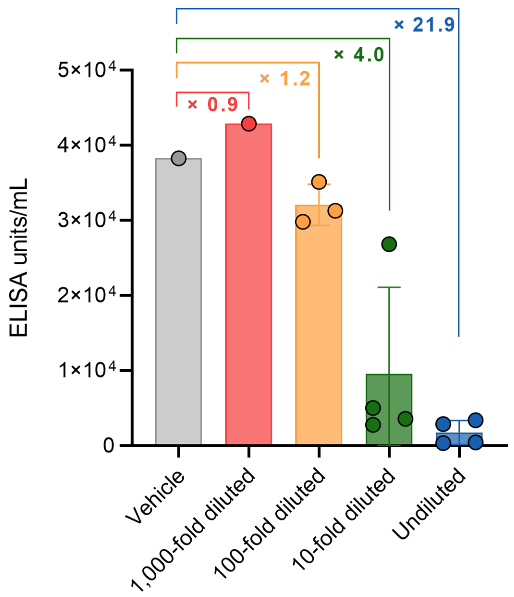

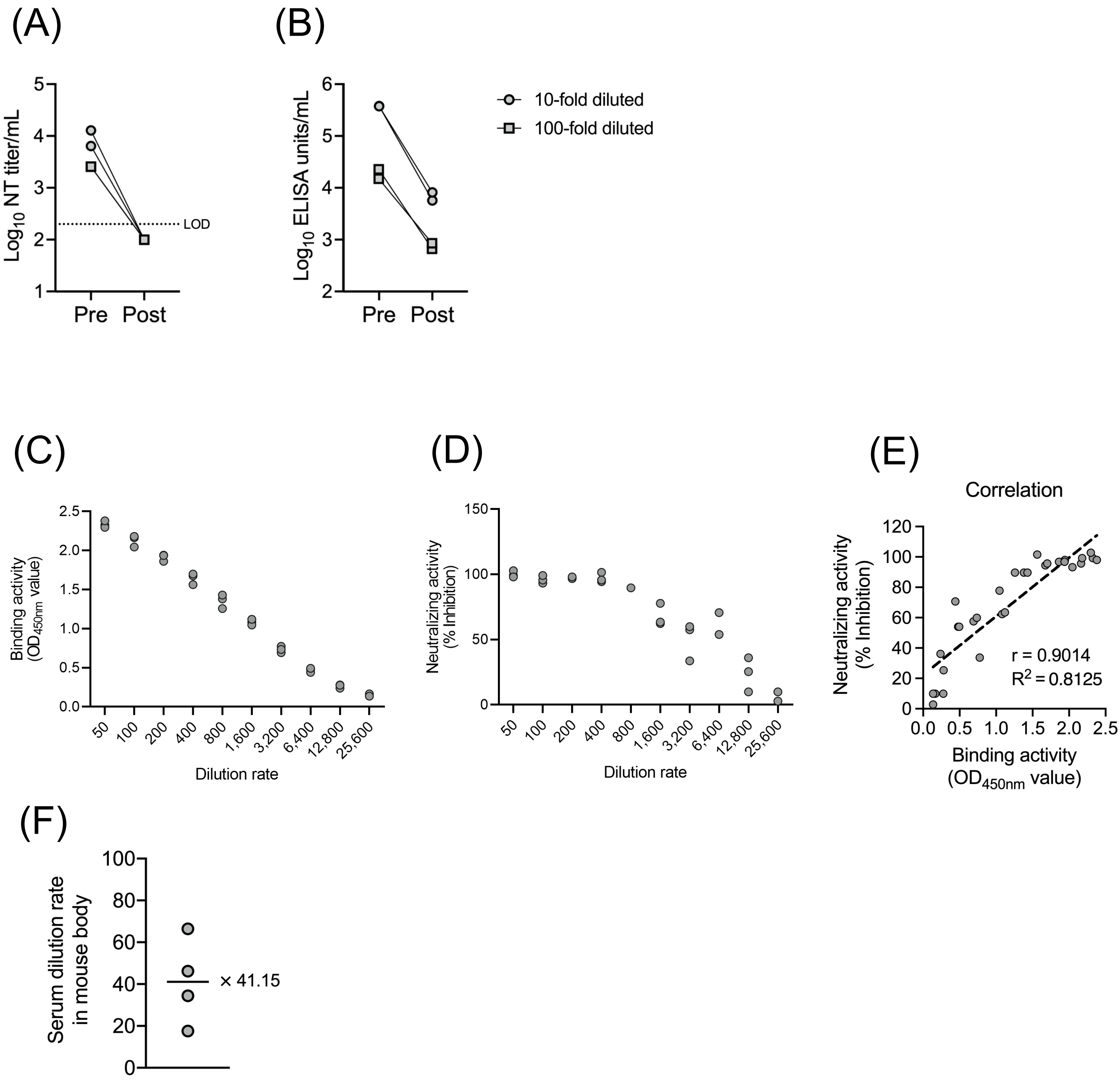

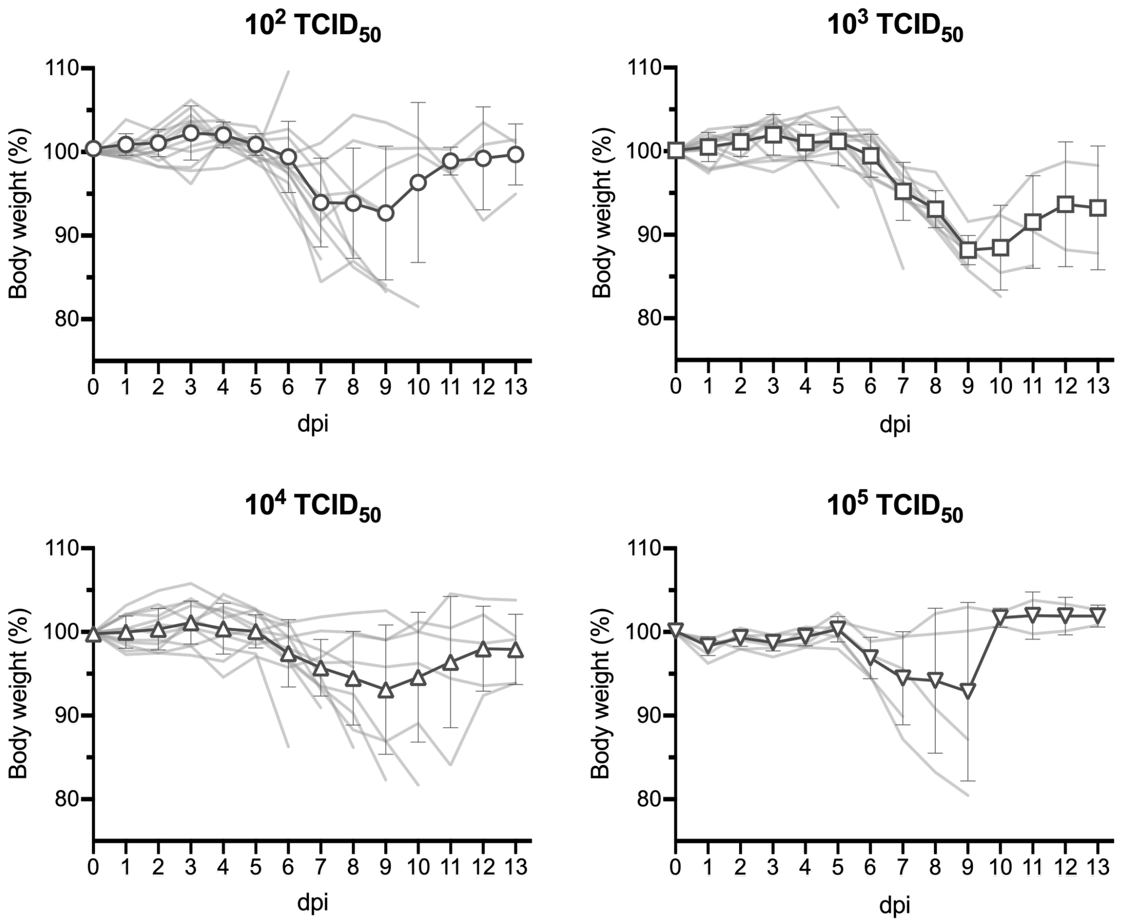

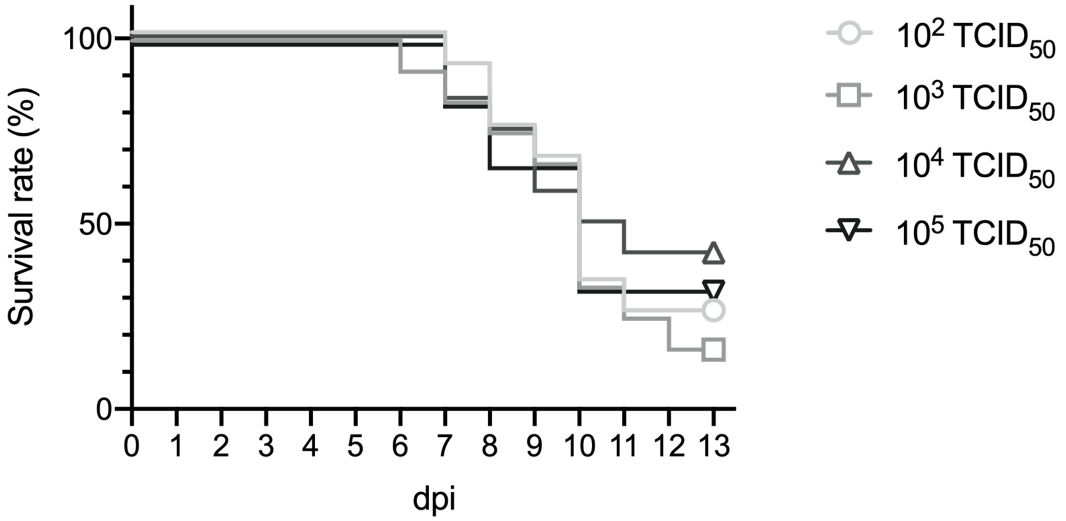

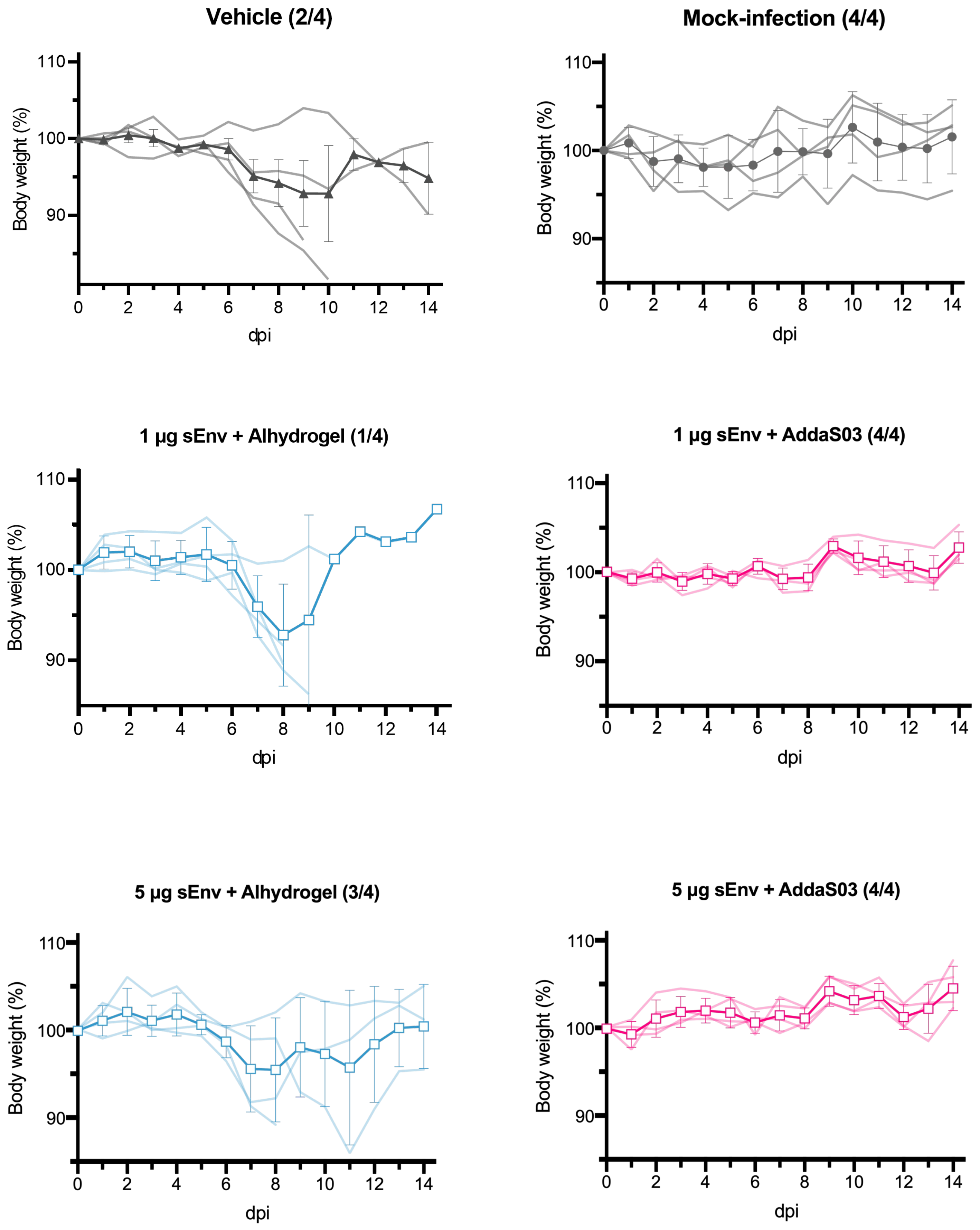
